# Leveraging Transcriptomics-Based Approaches to Enhance Genomic Prediction: Integrating SNPs and gene-networks for Cotton Fibre Quality Improvement

**DOI:** 10.1101/2024.02.14.580398

**Authors:** Nima Khalilisamani, Zitong Li, Filomena A. Pettolino, Philippe Moncuquet, Antonio Reverter, Colleen P. MacMillan

## Abstract

The cotton genome *(Gossypium hirsutum)* contains ∼ 80K protein-coding genes, making precision breeding for complex traits a challenge. This study tested biology-informed approaches to improve genomic prediction (GP) accuracy for cotton fibre traits to help accelerate precision breeding of valuable traits. The study’s foundational approach was the use of RNA-seq data from key time points during fibre development, namely fibre cells undergoing primary, transition, and secondary wall development. The test approaches included using a range of summary statistics from RNA-seq analysis such as gene Differential Expression (DE). The three test approaches included DE genes overall, target pairwise DE lists informed by gene functional annotation, and finally, gene-network-clusters created based on Partial Correlation and Information Theory (PCIT) as the prior information in Bayesian GP models. The most promising improvements in GP accuracy were at the level of ∼ 5% increase by using PCIT-based gene-network clusters as the prior knowledge network neighbours of key genes, and for the traits of cotton fibre Elongation and Strength. These results indicate that there is scope to help improve precision breeding of target traits by incorporating biology-based inference into GP models, and points to specific approaches to achieve this.

**KEY MESSAGE:** Improved Genomic Prediction accuracy of cotton fibre quality traits for helping accelerate precision breeding can be achieved by using biology-based prior knowledge and gene-network clusters.

## INTRODUCTION

Cotton (*Gossypium spp*.) is one of the most economically significant crops, providing fibre and oilseed products worldwide. In recent years, there has been an increasing demand for cotton with improved agronomic traits, which are crucial determinants of textile quality. Traditional phenotype-based breeding approaches have been limited by the time-consuming and labour-intensive nature of phenotypic evaluation and the underlying genetic complexity of major traits such as those spanning yield and quality. To help accelerate breeding, genomic prediction (GP) is a relatively recent strategy being researched for crop and tree breeding that utilizes genome-wide data to estimate breeding values. GP would enable genomic selection (GS) of valuable germplasm with the desirable traits early in the breeding process based on genetic information, and consequently help shorten the breeding cycle, reduce resource intensity, and facilitate precision of target traits. Furthermore, advancements in high-throughput sequencing technologies have raised many possibilities for the integration of omics-based approaches, particularly transcriptomics, to help accelerate the development of GS and GP strategies in cotton breeding.

Cotton seed fibre quality traits are important considerations in the global commodity market. These traits include fibre Length, Strength, and Elongation amongst others such as Micronaire, Short Fibre Index, Uniformity and Color. A key component of fibre quality is the cell wall whose composition plays a pivotal role in determining the fibre’s physical properties (Pettolino et al 2022). Although thousands of genes are expressed during fiber development (MacMillan & Birke et al 2017; Ma et al. 2018; Li et al. 2020; Gallagher et al. 2020) a majority of the key contributors to fibre traits remain unidentified. Known cell wall genes that affect the Strength and Elongation properties of the cell walls of a range of angiosperm fibres include fasciclin-like arabinogalactan proteins (FLAs) (MacMillan et al 2010; MacMillan et al. 2015; Ma et al. 2022), as well as cellulose synthases and cell wall biosynthetic enzymes that impact on fibre quality traits such as Length, Strength, and Elongation (Liang et al. 2015; Huang et al. 2015; Qin et al. 2011). Understanding the regulatory mechanisms of and the interactions among these genes is crucial for unravelling the genetic basis of these desirable traits.

Transcriptomics allows for the quantification of gene expression levels across different tissues, developmental stages, and environmental conditions. By utilizing differential expression (DE) analysis, researchers have identified the possibility of identifying genes that are differentially regulated in response to favourable traits thus using a biological basis for precision breeding approaches. This information, in turn, can guide the development of precise breeding strategies using biological understanding. Moreover, transcriptomic data have been providing valuable insights into the molecular mechanisms underlying these traits, aiding in the discovery of candidate genes and potential regulatory networks involved in their development (Zhang et al. 2015).

Gene-network analysis further enhances our understanding of the complex interactions between genes, proteins, and regulatory elements, and how they contribute to the expression of desirable traits in cotton. By constructing gene co-expression networks, it is now possible to identify highly interconnected gene-networks that may be functionally related and potentially associated with the phenotypic traits of interest, using computational graphical learning approaches such as Partial Correlation and Information theory (PCIT) (Reverter 2008; Watson-Haigh 2009). Genes within these networks can serve as key regulators or biomarkers of the traits, offering opportunities for targeted manipulation or marker-assisted selection (Xu et al. 2011; Deng et al. 2012; Gu et al. 2020).

Integrating omics-based data, in particular the transcriptome data, with GP approaches is becoming an increasingly popular strategy considered to hold potential for enhancing plant breeding (e.g. Hu et al. 2019; Azodi et al. 2020; Liu et al. 2020; Liu et al. 2022; Wang et al. 2023). Furthermore, Single Nucleotide Polymorphisms (SNPs) are commonly used as molecular markers in GS, capturing the genetic variation responsible for phenotypic variation. By incorporating transcriptomic information, which is more directly linked to phenotypes than SNPs, there could be potential to increase the accuracy of GP. One strategy for this is to measure the transcriptome data for the lines available for the GP study and then integrate the transcriptome and SNP data to jointly conduct prediction analysis (e.g. Hu et al. 2019; Perez et al. 2022).

Alternatively, differential expression (DE) analysis can be conducted using transcriptome data alone, and then the DE genes detected from the analysis can be used as knowledge in the GP model, e.g. by adding a specific weight to the effects of SNPs that are linked to the DE regions. Innovative approaches taken in this research outlined here include not only using DE genes, but also network analysis, and also considering biological relevant time points.

Biologically relevant transcriptomes can be important for identifying the DNA elements involved in a trait. For this study, a key element is the capture of synchronous single-cell transcriptomes across biologically relevant developmental stages (Fig.1A). The cotton seed-fibre is a single cell and its development follows a clear sequence, from seed fibre cell initiation just before the day of flowering, rapid cell growth to > 3 cm long with a soft primary cell wall over ∼2.5 weeks, a transition phase, then a secondary cell wall deposition phase during which cell-extension ceases and a thick secondary cell wall is deposited on the inner surface of the cell’s primary wall, a cell maturation and programmed cell death phase, and then finally yielding a dry mature fibre cell after ∼ 2 months that is harvested for trade on the global natural fibre commodity market.

**FIG.1.**
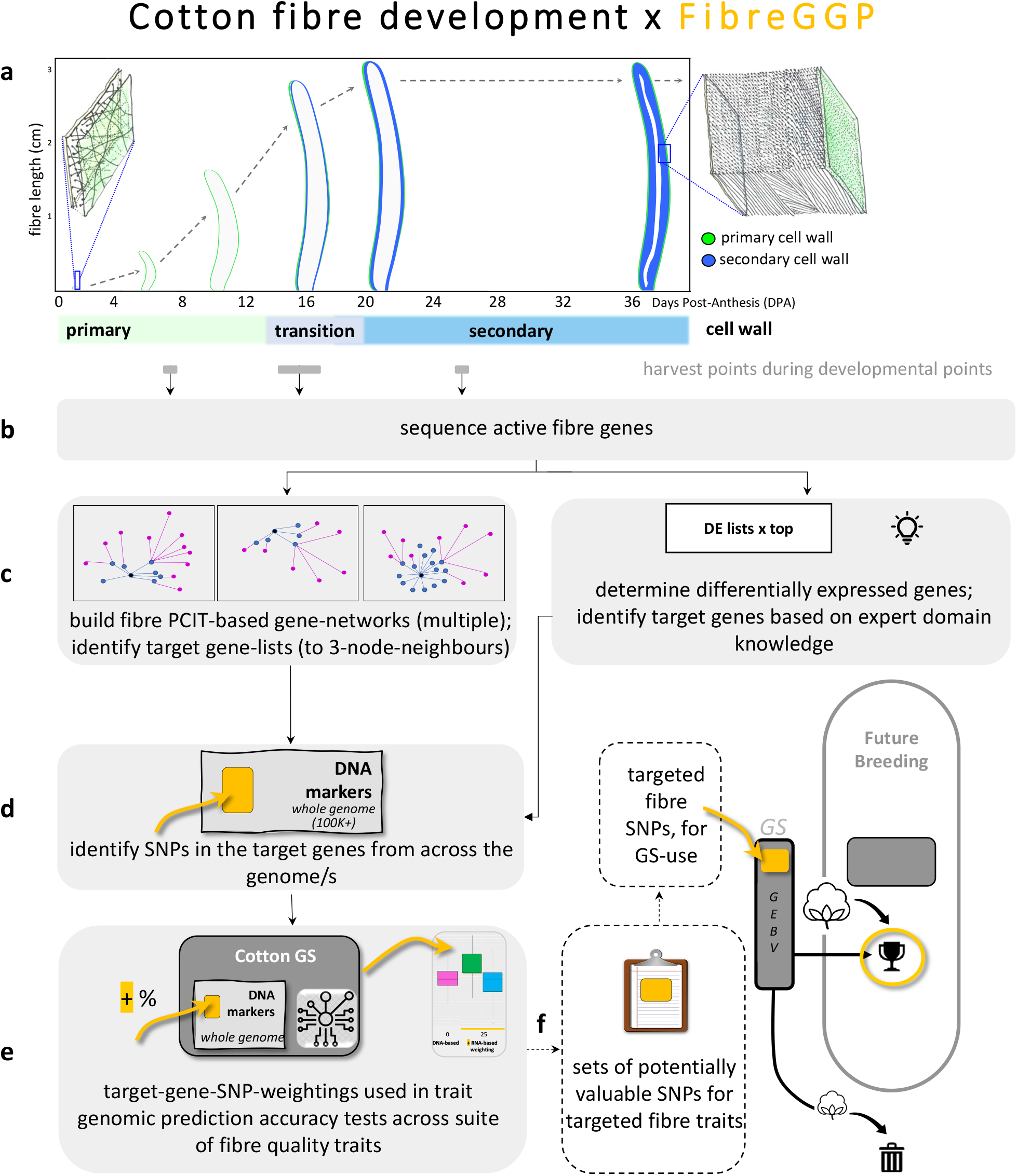
Cotton fibre development and the FibreGGP workflow undertaken in this study, and its links to cotton breeding. **(a)** Fibre genes expressed in key points of development are identified through targeted harvesting and RNA sequencing. A fibre cell emerges on the surface of the cotton seed (ovules) on the day of anthesis (flowering; i.e.,0 DPA); this fibre cell has a thin expanding primary cell wall. The fibre cell grows to several cm long over a period of 2-3 weeks. Towards the end of its growth phase, the fibre cell wall starts its transition to deposition of the secondary wall that continues for several weeks until a very thick secondary wall has been formed, and the cell matures, dehydrates, and dies. The mature fibre is typically harvested for commercial purposes once the cotton boll has fully opened ∼ 60+ DPA. Fibre expressed genes were obtained at three points in development (shown in grey bars) i.e., primary growth phase at 7DPA, a transition point at an average of 16DPA, and during secondary cell wall formation once fibre cell growth had largely ceased at 25DPA. **(b)** The fibre-expressed genes are sequenced via RNAseq. **(c)** Fibre-expressed genes are analysed in two ways to create target gene lists, via “PCIT gene-networks” i.e. that generate specialist gene-networks using PCIT and up to 3 node-neighbours in key gene-networks, and via “DE lists” i.e. differential gene-expression top-lists via expert domain knowledge. **(d)** The target gene lists networks guide the identification of SNPs based on the expressed fibre genes. **(e)** The fibre-guided SNPs are weighted in GS algorithms and tested for improved prediction accuracy of key fibre traits. **(f)** How the target SNPs are integrated into the breeding pipeline (dashed line; Not part of this study). Long- and short-lists of desirable target SNPs yielding positive results arepotentially fed into the cotton breeding pipeline. GEBV = Genetic Estimated Breeding Values.

In this study, we aimed to investigate the potential of integrating transcriptomic information obtained through RNA-Seq analysis as a pre-knowledge in GP models in cotton breeding. By leveraging developmental time course differential expression and gene-network analysis, we identify candidate genes and gene modules associated with fibre Length, Elongation, and Strength. Next, these genes and gene modules were considered as prior information in a well-known Bayesian GP model (i.e. Bayes C model), where a small number of SNPs located within or near to the relevant gene regions are assigned with a different prior which weighted their genetic effects more heavily than others. We then evaluate whether this weighted Bayesian model would provide enhanced prediction accuracy compared to the standard Bayes C model, on a previously published cotton genome selection data set collected from the CSIRO breeding program (Li et al. 2022). This research has the potential to advance cotton breeding strategies by providing a more comprehensive understanding of the genetic architecture underlying desirable traits and enabling the identification of superior genotypes at an earlier stage.

The foundation basis of the workflow undertaken in this study is the biology of the cotton seed fibre. This framework, “<u>Fibre G</u>ene-network Guided <u>G</u>enomic <u>P</u>rediction” (FibreGGP), is outlined in Fig.1. Gene expression at specific fibre developmental points was determined with RNA extracted at key points in the primary, transition, and secondary cell wall stages of the cell’s development (Fig.1a). In this study, we interrogate the expression of all the genes expressed at each of these developmental points (Fig.1b). To examine potential improvements in GP accuracy of cotton fibre quality traits, a Bayesian regression approach using various summary statistics derived from transcriptome data analysis as prior information was tested in targeted genes from the cotton genome (Fig.1c – e) across three scenarios. Scenario 1 tested all the DE genes arising from pairwise comparisons. Scenario 2 tested select DE genes that were considered biologically relevant to the trait of interest. Scenario 3 tested a highly targeted set of DE genes derived from PCIT-based gene-networks with particular key genes, and their 1st, 2nd and 3rd network-neighbours. In this scenario, we tested key genes linked to the biomechanical properties of strength and elasticity of angiosperm-fibre secondary cell walls i.e. a sub-class of FLAs. Ultimately our framework would potentially be used to enhance the precision-breeding of cotton for select traits (Fig.1f).

## MATERIALS AND METHODS

### Transcriptomics, Genomics, and Phenomics Data

The transcriptome data used in this study were retrieved from MacMillan & Birke et al. (2017). The authors collected three replicates for each of the three stages of fibre development i.e. at days 7, 16, and 25 Days Post Anthesis (DPA). Genomic and phenotypic data were obtained from Li et al. (2022), representing 1907 samples collected from 1994 to 2017 and 12296 informative SNPs. Our focus was three fibre quality traits i.e. fibre Length (upper half-mean length of sample), Strength (the breaking point force of a bundle of fibres of a given weight and fineness, g tex−1), and Elongation (fibre bundle extension force up to its breaking point, expressed as a % increase over its original length).

### RNA-seq, PCIT, and Gene-Network Analyses

RNA-seq analysis was conducted on the raw transcriptome data (above). FastQC software (Andrews 2010) was employed to assess the quality of the RNA-seq data. The genome of *G. hirsutum* version 1 (Li et al. 2015; Zhang et al. 2015) was indexed using HISAT2 (Kim et al. 2015) to facilitate the sequence alignment. Afterwards, HISAT2 was used to align the RNA-seq files to the reference genome, generating Sequence Alignment/Map (.sam) file format. The .sam files were converted to Binary Alignment/Map (.bam) files using SAMtools (Kim et al. 2015) for further analysis. The mapped files were merged using Cuffmerge (Trapnell et al. 2012), and the resulting merged map file was compared to the reference genome using Cuffcompare (Trapnell et al. 2012). The Gossypium hirsutum v3.1, DOE-JGI, http://phytozome.jgi.doe.gov was used as the Reference Genome. Next, Stringtie (Pertea et al. 2015) was used to estimate the abundance of genes/transcripts in the mapped file. This step resulted in the construction of a gene count matrix for differential expression analysis (DE). Fragments Per Kilobase of transcript per Million mapped reads (FPKM) was used to measure and normalize expression levels in the analyses. Consequently, the DE analysis was performed using the DESeq2 package (Love et al. 2014), with DE genes identified using the false discovery rate (FDR) threshold of 0.001 and log2foldchange of 1.5. The differential expression analysis was performed for comparison of three stages of fibre e.g., fibre 07 vs 16, fibre 07 vs 25 and fibre 16 vs 25.

After differential expression analysis, PCIT (Reverter et al 2008; Watson-Haigh et al. 2009) was used to estimate the pair-wise correlation between DE genes, and consequently correlation-based gene-networks were generated using Cytoscape 3.9.1. The gene-networks were used to identify clusters of genes potentially linked to specific fibre quality traits. Each gene-network was interrogated by identifying a key fibre-trait gene and then identifying its 1^st^, 2^nd^, and 3^rd^ neighbours of genes in the network associated with fibre-quality traits.

### Identification of SNPs in target genes / gene regions

Various scenarios were used in the identification of SNPs that were subsequently tested for GP accuracy of various traits. SNPs for various traits were identified based on DE gene lists, expert domain knowledge, and via PCIT-based gene-networks.

*Scenario 1:* DE genes were obtained via DESeq2 from three fibre developmental stages comparing fibre 07 vs 16, fibre 07 vs 25 and fibre 16 vs 25. Genome coordinates of the DE genes were identified, followed by identification of SNPs within the DE genes.

*Scenario 2:* Annotation i.e. biological information from the DE gene lists was used to identify a subset of SNPs associated with Length and Strength. Expert domain knowledge of fibre quality traits was employed to identify target genes for these traits e.g. down regulated genes from the fibre 07 vs 16 DE gene data set that were considered to be either involved in Length, or upregulated genes from fibre 07 vs 25 DE gene set for Strength. In addition, within each data set of this scenario, SNPs were identified either in the exact gene region, within a 1 kb region, and within a 10 kb region. This resulted in having 3 sets of SNPs for each trait where applicable except for Length in which did not have SNPs in 1 kb of DE genes. Using this scenario there were 7 sub scenarios: L0, L1, L3, S0, S1, S2, S3 were number 0 represents unweighted SNP scheme and numbers 1, 2 and 3 indicate the presence of SNPs in the exact gene location, within 1 kb, and within 10 kb of the selected DE genes respectively. L represents Length and S represents Strength.

*Scenario 3:* Three sets of DE genes from fibre 07 vs 25 were identified as crucial for fibre quality namely FLA7, FLA11 and FLA12 (this was based on known genes closely involved in the biomechanics of plant fibres, MacMillan et al. 2010, 2015). Next, each of these three sets of genes was extended to include their 1^st^, 2^nd^ and 3^rd^ network neighbours, and then relevant SNPs were identified before analysis for impact on Strength, Length, and Elongation.

### Bayesian GP model accounting for SNPs associated with DE gene regions

In a GP model, we aimed to add more on the effects of subset of SNPs linked to DE gene regions or their close neighbours in gene-networks, which can be done by using specific priors for those SNPs. This idea is illustrated on the well-known Bayes C linear regression model, with its likelihood model form defined as follows:

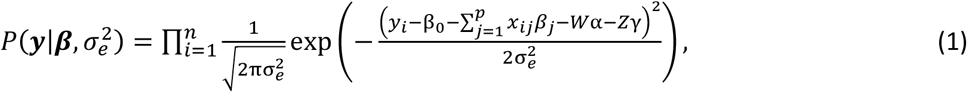

where *y*_*i*_ is the phenotype record of the *i*th individual (*i*=1,…,*n*; *n* is the total number of individuals), *β*_0_ is the model intercept, *e*_*i*_ is the residual error: 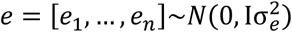(mutually independent for 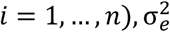 is the residual variance, *β*_*j*_ (*j*=1, …, *p*) is the regression coefficient representing the additive genetic effect of the marker *j*. The genetic effect *β*_*j*_ was assigned with a spike and slab prior (Ishwaran and Rao 2005) to the regression parameters as follows:

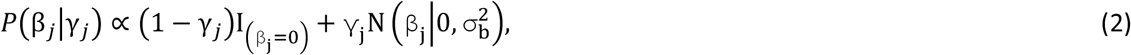

where γ_*j*_ is a binary indicator variable to tell whether the genetic effect of SNP *j* should be non-negligible and follow a normal distribution, or whether the effect is small and assigned with a zero value. In the standard Bayes C model, the indicator variable γ_*j*_ and the variance component 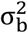 are further assigned with priors of Bernoulli: *Bern*(γ_*j*∣_ π) and Inverse chi-squared: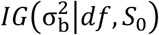, respectively. In the Inverse gamma prior 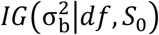, the parameters df=5 and *S*_0_ = var(*y*) × *R*^2^ × (*df* + 2)/*MSx*, with R^2^=0.5, assuming that 50% of the phenotype variance is explained by the whole set of SNPs. In the Bernoulli prior *Bern*(γ_*j*∣_ π), the parameter π was further assigned with a Beta prior Beta (π|p_0_, π_0_), with p_0_ =50, and π_0_ =0.5. The spike and slab prior (13) are often referred as the Bayes C model (Habier et al. 2011) in the GP literature.

To add more weights on the effects of SNPs linked to DE gene regions, an alternative prior to (2) was specified:

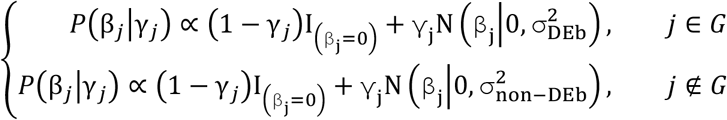

where G represents a specific subset of SNPs linked to a DE gene region or their neighbours in a gene-network, separate variances 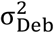 and 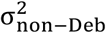 were given to a SNP depending on whether it was presented in G. Those variances were then assigned with different hyper-priors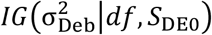, and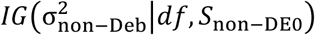,respectively, where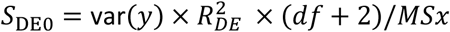, and 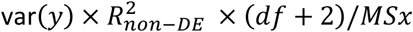.the choice of 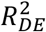 and 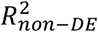 determined how much weights were assumed for SNPs within G. We used the following three combinations: (i) 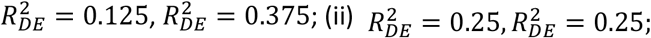; and 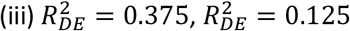. These settings correspond to the SNPs within G assumed to explain 25%, 50% and 75% of the total genetic variance, respectively. SNPs identified in the DE genes were weighted at either low (25%), medium (50%) or high (75%) levels in the GP calculations to test whether their role in the target traits were significant for improving GP accuracy. The improved prediction accuracy would indicate that these SNPs linked to DE genes or gene networks play a more important role than other SNPs, since they were weighted more in the Bayesian regression model to predict fibre qualities.

The Bayesian C algorithm utilizes the Markov Chain Monte Carlo (MCMC) method to sample from the joint posterior distribution of the model parameters. MCMC generates a sequence of samples that asymptotically approximates the true posterior distribution. It allows for uncertainty quantification and inference based on the posterior samples. In practice, 100 00 samples were generated as burn-in, and the rest 200 00 samples were thinned to keep every 20^th^ samples to reduce serial correlation. Hence, 1000 samples were collected to approximate the posterior.

We took the latest 334 new lines collected during 2017/18 season as the test population, and the rest 1051 lines collected prior to 2017 as the training population (Li et al. 2022). The Pearson correlation between the genomic estimated and true phenotypes of the test population was defined as prediction accuracy. The prediction method was implemented repeated 60 times, and the average prediction accuracy was calculated to reduce the randomness introduced by the MCMC sampling.

## RESULTS

### DE results from key fibre RNAseq analysis provided a large target gene list and a large range of SNPs

Across the three key developmental stages of Coker315-11 seed fibre, the pairwise DE analyses found that there were about three to eight thousand differentially expressed genes (at a stringent P-adjusted value <0.001) (Supplementary 1). These findings are within the ballpark reported in detail regarding gene classes and pathways using the *G. raimondii* (MacMillan & Birke et al. 2017) and also found here but instead using the more recent *G. hirsutum* reference genome. The gene classes span transcription factors, cell wall biosynthetic genes, multiple metabolic and structural genes, and unknown function genes. The Fibre7DPA vs Fibre16DPA comparison (primary vs transition wall stage comparison) identified 2928 differentially expressed genes, with 1487 upregulated and 1441 down-regulated genes, with log fold-changes ranging from 13 to 2. The Fibre16DPA vs Fibre25DPA (transition vs secondary wall stage comparison) comparison identified 3917 differentially expressed genes, with 1956 upregulated and 1961 down-regulated, with log fold-changes ranging from 16 to 2. The Fibre7DPA vs Fibre25DPA (primary vs secondary wall stage comparison) comparison identified 7678 differentially expressed genes, with 3931 upregulated and 3747 down-regulated, with log fold-changes ranging from 15 to 2.

The Coker fibre DE gene lists enabled the identification of a suite of SNPs across the 63K SNP chip array of Hulse-Kemp et al. 2015, and these SNPs ranged in number across DE gene regions (Supplementary 2). For the DE genes from the Fibre7DPA vs Fibre16DPA comparison (primary vs transition wall stage comparison) 85 SNPs were found within the gene coding regions, 54 SNPs in the 1kb gene-coding flanking regions, and 441 SNPs in the 10kb gene-coding flanking regions. For the DE genes from the Fibre16DPA vs Fibre25DPA comparison (transition vs secondary wall stage comparison) 87 SNPs were found within the gene coding regions, 72 SNPs in the 1kb gene-coding flanking regions, and 594 SNPs in the 10kb gene-coding flanking regions. For the DE genes from the Fibre7DPA vs Fibre25DPA comparison (primary vs secondary wall stage comparison) 260 SNPs were found within the gene coding regions, 135 SNPs in the 1kb gene-coding flanking regions, and 1183 SNPs in the 10kb gene-coding flanking regions.

### Weighted-SNPs from DE genes alone did not increase GP accuracy for fibre quality traits (Scenario 1)

The first scenario tested was weighting of DE genes alone (Fig.1b). The result of weighting SNPs from the DE genes for three fibre development stages (fibre 07 vs 16, fibre 07 vs 25 and fibre 16 vs 25) for estimating GP accuracy is presented in Fig.2; the summary results of the prediction accuracies are presented in Supplementary 3. Adding more weights on SNPs linked to DE genes in the Bayesian GP model were largely ineffective e.g. accuracy of predictions showed little or no improvement for Elongation (Fig.2a) and Strength (Fig.2c) but a reduction in accuracy was noticeable for Length (Fig.2b) as compared to the scenario where no weight was considered. Furthermore, the results were largely distributed, representing outliers, which might infer that the results were not reliable. On the other hand, in a few instances weighting of SNPs at different levels (25%, 50%, 75%) improved GP accuracy above the base-level. For example, for Elongation increased GP accuracy was found with 50% and 75% SNP weighting of all DE gene lists. This did not extend to Length or Strength.

**FIG.2.**
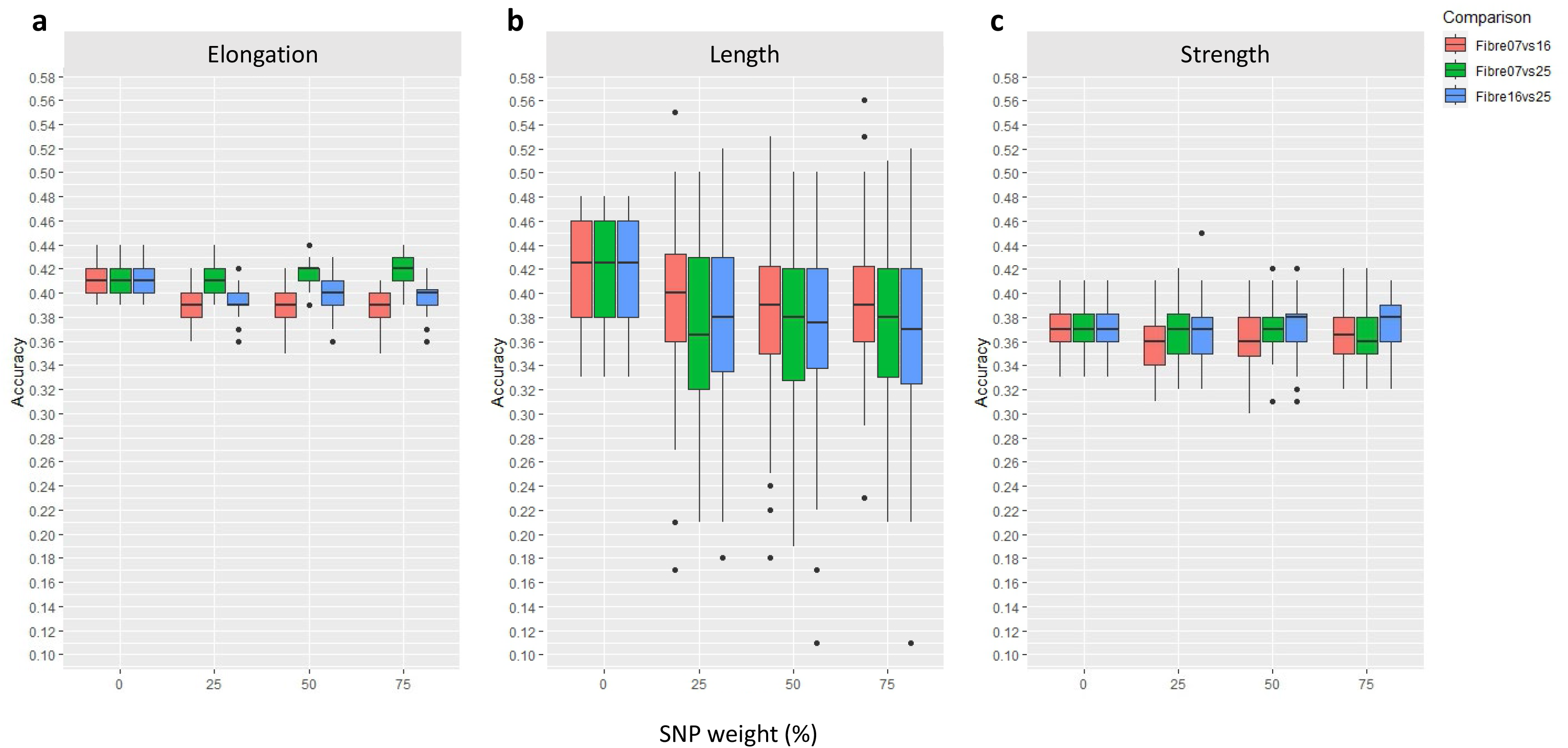
Accuracy of genomic predictions for fibre quality traits of **(a)** Elongation **(b)** Length and **(c)** Strength using SNPs weighted based on their coordinates compared to DE genes across three different fibre developmental stages (fibre 07 vs 16, fibre 07 vs 25 and fibre 16 vs 25). The results were presented for SNP weights of 0, 25, 50 and 75 percent. The legends are showing Fibre 07 vs 16, Fibre 07 vs 25 and Fibre 16 vs 25 in red, green and blue respectively. Each box shows 50% of the data-range. A Horizontal bar within each box shows the median. Vertical bars extending from each box shows the range of the remaining data. Dots indicate outliers.

### Weighted-SNPs based on the annotation of targeted DE genes increased GP accuracy (Scenario 2)

Accuracy of GP using DE genes in conjunction with their annotation was a specific scenario tested in our workflow (Fig.1c); the results are shown in Fig.3. The target traits were Length and Strength, and the DE genes targeted include those down-regulated at Fibre 07 vs 16 and up-regulated Fibre 07 vs 25. The results showed that using annotation for weighting SNPs marginally increased the accuracy of prediction for Strength. For instance, when SNPs in exact, within 1kb and 10 kb of up-regulated DE genes from fibre 07 vs 25 were used, a 5.4% increase in accuracy of Strength was found (Fig.3B; S3, 75% weighting). On the other hand, for Length, a marginal GP increase was seen only when SNPs within and up to 1kb of the target DE genes were used, otherwise a downward shift was found if SNPs within 10kb were used (Fig.3A). A summary statistic of accuracies and number of weighted SNPs in each sub-scenario is provided in Supplementary 4.

**FIG.3.**
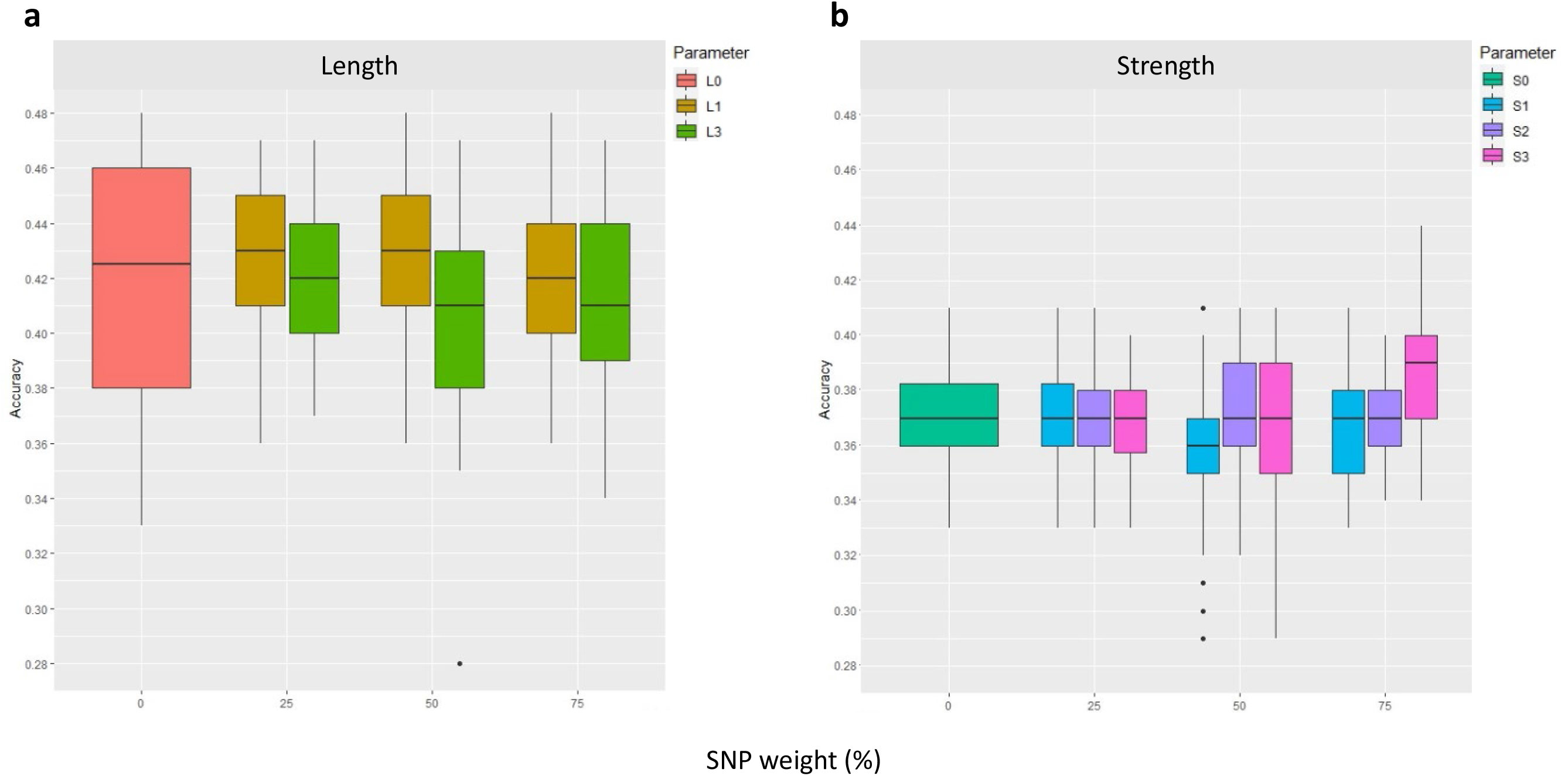
GP accuracy for fibre quality traits of **(a)** Length and **(b)** Strength, using a scenario-based approach. In this scenario, SNPs were weighted based on their coordinates in relation to select DE genes. The DE genes were from fibre developmental points of fibre 07 vs 16 for Strength and fibre 07 vs 25 for Length. The results were presented for SNP weights of 0, 25, 50 and 75 percent. The legends show L and S for Length and Strength in order. The numbers 0, 1, 2, 3 indicate (0) the unweighted scenario, and where SNPs were available in (1) exact; (2) exact and within 1 kb; (3) exact, within 1 kb and within 10 kb; of DE genes associated with down and up-regulated data sets of fibre developmental stages. No unique SNPs were found in the region L2, and hence no results reported.

### PCIT gene-network-clusters for SNP-weighting improved GP accuracy across traits, particularly with up to 3^rd^-network neighbours (Scenario 3)

Highly targeted gene-lists for trait-SNP identification were generated using PCIT-based gene-networks, and these included a nested gene-network-neighbour approach (Fig.1c). The PCIT-based gene-network was filtered for gene-network neighbours of the target FLA genes (FLA7, FLA11, FLA12), across their 1^st^, 2^nd^, and 3^rd^ network-neighbours; on average for each FLA tested about 300 (1^st^), 1200 (2^nd^) and 1900 (3^rd^) neighbours were identified in each gene-cluster respectively (Supplementary 5). In each case, the gene-network neighbours included an intriguing range of transcription factors, cell wall biosynthetic genes, other genes of known and unknown function, as well as other FLAs (Supplementary 5, individual lists).

Improved accuracy of GPs with up to the 3^rd^ PCIT-based gene-network neighbours was found, across the FLA7, FLA11 and FLA12 genes with the fibre 16 vs 25 DE lists (Fig.4). This increase in GP accuracy was seen for Elongation and Strength as compared to unweighted scenarios. The use of PCIT gene-network 1^st^ and 2^nd^ neighbours improved GP by 4.8% over that for unweighted GP for fibre Elongation in the case of FLA7 and FLA11, and inclusion of 3^rd^ network neighbours had no further effects, either negative or positive.

**FIG.4.**
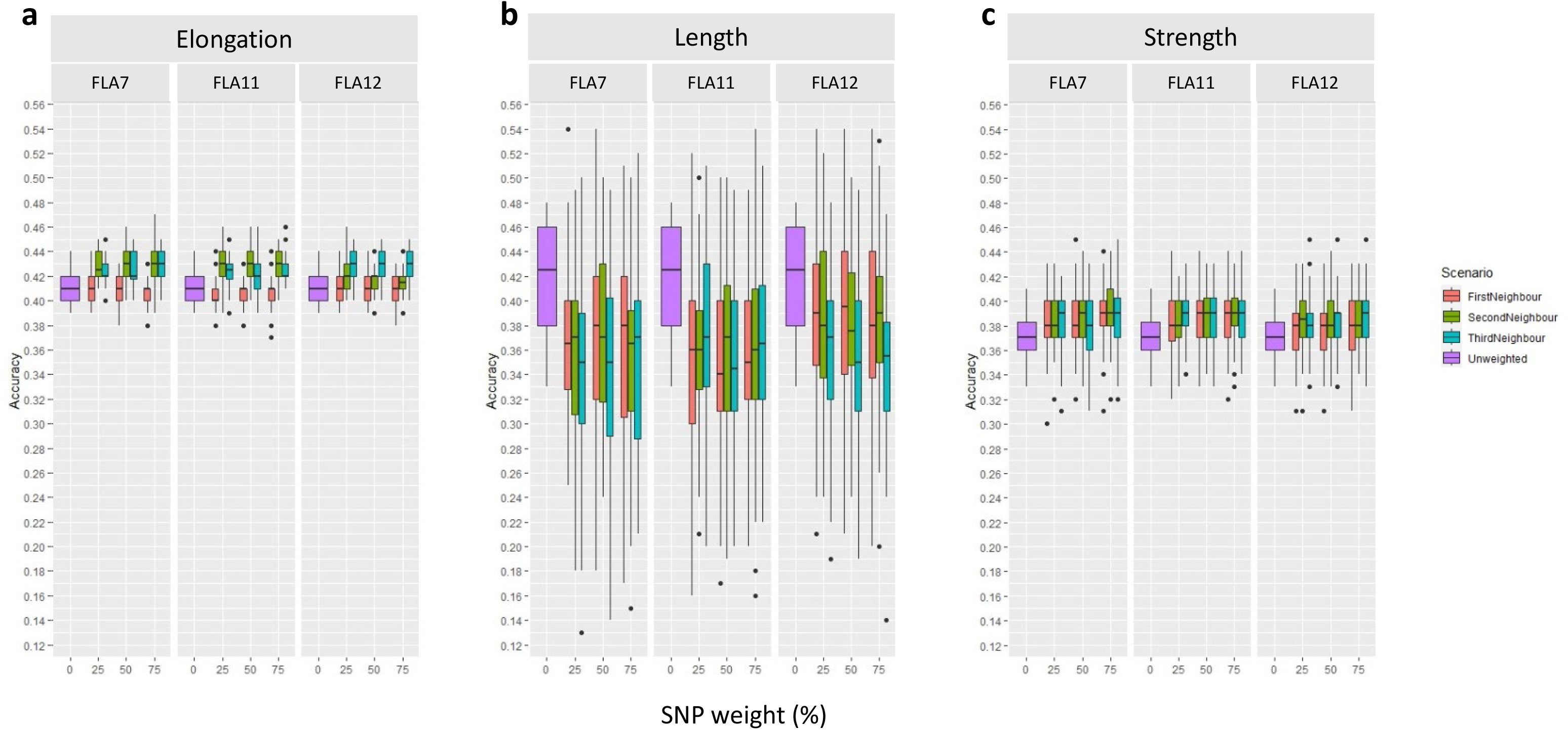
GP accuracy for fibre quality traits of **(a)** Elongation, **(b)** Length and **(c)** Strength using PCIT, gene-networks and weighted SNPs (Scenario 3). In this scenario, PCIT and gene-networks were used with a focus on key genes of FLA7, FLA11 and FLA12, and the first, second and third neighbours of each gene in the gene-network-clusters. SNPs were weighted based on their coordinates in the selected DE genes in the network clusters from the fibre 16 vs 25 comparison. The results were presented for SNP weighted at 0, 25, 50 and 75 percent.

Inclusion of 3^rd^ network neighbours increased Elongation GP accuracy in the case of FLA12. Weighting of the highly targeted SNPs at either 25%, 50%, or 75% did not provide additional GP accuracy for Elongation over that seen with the up to 3^rd^-network-neighbour approach. Increased accuracy of GP was also found for Strength, and in this instance up to 5.1%, when up to the 3^rd^ network neighbours were included. This was the case for FLA7, FLA11, and FLA12 gene-networks. However, use of the FLA7, FLA11 and FLA12 gene-network clusters did not improve GP for Length, and in fact, reduced GP compared to the unweighted GP. As these FLA genes are not known to be involved in Length, this result was somewhat expected; also, the developmental time points used in this scenario (F 16 v 25) are beyond when most of the fibre’s growth to its final length has occurred. In comparison, Scenario 1 and 2 only lead to 2-3% increase of prediction accuracy, but Scenario 3 lead to 5% increase.

## DISCUSSION

This research explored the potential integration of transcriptomic information derived from RNA-Seq analysis into GP models for cotton breeding (Fig.1). The study focused on elucidating key genes and gene-networks associated with essential fibre traits—such as Length, Elongation, and Strength—to inform more accurate GP. Leveraging DE analysis across three crucial developmental stages of seed fibre, numerous genes were identified, revealing thousands DE genes across these stages. These genes span various classes, including transcription factors, cell wall biosynthetic genes, and structural genes, shedding light on their crucial roles in fibre development. Integration of this transcriptomic data with genomic information allowed for the identification of SNPs located within or near specific DE genes. However, attempts to directly add more weight on these SNPs in the GP model based on DE genes or their annotations did not notably improve GP accuracy. In contrast, when incorporating information from PCIT-based gene-networks derived from DE genes and their 1^st^, 2^nd^ and 3^rd^ neighbouring nodes, the GP model exhibited enhanced accuracy for predicting Elongation and Strength. In other words, use of Scenario 3 lead to ∼ 5% increases, whereas Scenarios 1 and 2 lead to 2-3 % increases. The accuracy for predicting fibre Length showed a reduction, possibly owing to the nuances of the chosen developmental stage comparison and the genes used to generate the network clusters. This comprehensive approach offers insights into potential candidate genes and networks that impact cotton fibre traits, laying the groundwork for more refined and accurate GS strategies in cotton breeding.

PCIT-based gene-network clusters were effective in improving GP accuracy for cotton fibre quality traits. This was evident with a 4.8% improvement for Elongation and a 5.2% for Strength GP accuracy for cotton fibre quality. Inclusion of both the PCIT and the target gene-network approaches together proved valuable and opens up opportunities for advancing precision-breeding in cotton using such an approach. PCIT as an approach has also been of value in studies of animals and humans, with key regulator genes being identified as predictors controlling ∼500 genes associated with heifer puberty (Nguyen et al. 2018), significant suites of bovine fertility genes identified such as regulatory and functional genes as well as non-coding RNAs (Fonseca et al. 2022), pig gut microbiotal changes across intestinal tracts and their association with energy homeostasis (Crespo-Piazuelo et al. 2018), and in identifying key target genes in studies of human cancers (Li et al. 2022), asthma (Banerjee et al. 2021), and the SARS-CoV-2/human interactome (Guzzi et al. 2020).

The targeted gene-network-cluster scenario (PCIT-based) improved GP accuracy best with up to 3^rd^ neighbours, and flags potential utility in cotton breeding. Such a gene-network-neighbour clustering approach for GP in crop breeding is to our knowledge not currently being used, but our findings provide practical evidence, and workflows, to help achieve the vision and aspiration for future breeding such as large gene-to-phenotype networks that others have pointed to (e.g. Powell et al. 2022). With only three traits tested here across the various scenarios, our findings open up the possibility for further traits that could be more accurately predicted for cotton fibre and other traits. It also points to how large-scale target SNP populations can be identified for precision breeding.

Using a DE gene-only approach was not effective in improving GP accuracy in this study even with varying the extent of SNP weighting; this may point to the complexity of genome to phenome in crop development and target traits such as in cotton. In terms of DE gene studies for cotton fibre improvement, over the last 5 years dozens of articles have been published internationally, and generally these report substantial sets of DE genes with potential for impacting fibre quality. For example, these include studies combining GWAS and eQTLs to identify fibre cell wall development regulatory networks (Li et al. 2020), co-expression analyses for genes differentially expressed with high cotton fibre quality (Zou et al. 2019), and high-density genetic map studies for QTLs and genes for fibre quality and yield in cotton analysis of vegetative to reproductive transition and branching genes linked to planting density (Gu et al. 2020), and also linked to natural color formation in fibres (Tang et al. 2021). In our study, weighting SNPs increased their efficacy for GP accuracy depending on their location and this could be due to the fact that negative and positive SNPs can co-occur across chromosomal regions e.g. SNPs in enhancers, suppressor regions, promoters, etc.; when such SNPs are in key genes are important for a trait then the interplay of positive and negative SNPs could come into effect. Given the results of this study, such DE lists likely hold a rich wealth of important insights into the genes underlying the biology of fibre formation and use in GP accuracy for traits of interest.

Our strategy can be easily used in breeding practice. The transcriptome experiments and data analysis were done independently from the GP, and the outcome of transcriptome data analysis was used as prior information in Bayesian prediction models applicable for any cotton GP data set that targeted fibre qualities. This is much simpler and cost-effective to be applied in a breeding program compared to existing omics approaches which rely on measuring both the transcriptome and genomic data for thousands of samples (e.g. Hu et al. 2019; Azodi et al. 2020). As a future direction, evaluation will be required to determine whether this strategy can also be beneficial to the multiple environment GP analysis (i.e. prediction models with G×E interactions), which is essential from the application point of view in a plant breeding program.

In conclusion, the integration of omics-based approaches, particularly transcriptomics to guide SNP weighting, holds promise for enhancing GS in cotton breeding. By leveraging RNA-Seq data, DE analysis, PCIT-based gene-network analysis, and weighted SNP approaches, we can gain valuable insights into the genetic basis of important fibre traits. This knowledge could accelerate the development of improved cotton varieties with enhanced fibre Length, Elongation, and Strength, meeting the demands of the textile industry and ensuring the sustainability of cotton production.

## STATEMENTS AND DECLARATIONS

### Author Contributions

Conceptualization: CM FP; Data curation NK ZL CM; Formal Analysis NK CM FP; Funding acquisition CM FP; Investigation NK CM FP; Methodology CM FP NK ZL TR; Project administration CM FP; Resources AR PM ZL; Software AR NK ZL; Supervision CM FP; Visualization NK CM; Writing – original draft NK ZL CM; Writing – review & editing NK ZL FP PM AR CM

### Data Availability

The phenotype and genotype data were initially published in Li et al. (2022), and the data are publicly available at the CSIRO Data Access Portal. The transcriptome data, as well as workflows, will also be available at the CSIRO Data Access Portal upon the acceptance of the manuscript.

## Acknowledgements

The authors thank the teams at CSIRO Agriculture and Food, Cotton Breeding Australia, Cotton Seed Distributors for their ongoing support.

## SUPPLEMENTARY INFORMATION

**Supplementary file1**

CokerFibreDevelopmentDEPairwiseLists

**Supplementary file2**

CokerFibreDEgenes_SNPcoordinates

**Supplementary file3**

Accuracy prediction estimates for three fibre life stage comparisons

**Supplementary file4**

Summary statistic of accuracy of genomic predictions for fibre quality traits

**Supplementary file5**

CokerPCITbasedGeneNetworks GP and Neighbours

